# Adolescent rats show estrous cycle-mediated sex-differences in extinction of conditioned fear

**DOI:** 10.1101/2020.03.31.019273

**Authors:** Christina J Perry, Despina E Ganella, Ly Dao Nguyen, Xin Du, Katherine D Drummond, Sarah Whittle, Terence Y Pang, Jee Hyun Kim

## Abstract

Anxiety disorders are more prevalent in females than males, and frequently emerge during adolescence. Despite this, preclinical research commonly focuses on adult males. Here we use Pavlovian fear conditioning and extinction in adolescent male and female rats to understand sex- and age-dependent processes relevant to anxiety disorders. In experiment 1, 35-day-old male and female rats were exposed to 6 pairings of a conditioned stimulus (CS, a tone) with an aversive unconditioned stimulus (US, a footshock). The next day they were extinguished in a contextually distinct chamber, via 60 presentations of the CS without the US. Extinction recall was tested 24 hours later in the extinction context. Estrous phase was monitored by cytology on vaginal smears taken 1 hour after each behavioral session. In experiment 2, male and female rats were given sham surgery or gonadectomy at 21 days of age. They were then trained and tested as for experiment 1. We observed that females in proestrus or met/diestrus during extinction showed delayed extinction and impaired extinction recall the next day compared to males. Ovariectomy enhanced extinction for female rats, but orchidectomy delayed extinction for males. Plasma analyses showed that met/di/proestrus phases were associated with high estradiol levels. These findings suggest that high plasma estradiol levels impair extinction for adolescent females. While these results contradict what is observed in adult animals, they are consistent with the prevalence of anxiety disorders observed in females. Our findings have important implications for understanding and treating anxiety in adolescents, particularly where treatment involves extinction-based therapies.

## 1. Introduction

The prevalence of anxiety disorders is sex-dependent, with an almost two-fold increase in likelihood for females compared to males (Baxter et al., 2012). Prevalence is also dependent on age. Approximately three quarters of anxiety disorders are diagnosed during adolescence (Kessler et al., 2007), and adolescent onset is associated with a more severe long-term prognosis (Kim and Ganella, 2015; Newman et al., 1996). However, much preclinical research into anxiety disorders is restricted to adult males, neglecting the most affected group of adolescent females.

As a preclinical model, Pavlovian fear conditioning and extinction captures important cognitive aspects of anxiety disorder and post-traumatic stress disorder, including augmented retrieval of learned fear and the inability to inhibit fear in safe circumstances (Maren et al., 2013; Flores, Fullana, Soriano-Mas, & Andero, 2018). In this paradigm, subjects typically learn to fear a conditioned stimulus (CS) through repeated pairing of an aversive shock (the unconditioned stimulus, or US) with the CS. Extinction occurs when the subject is exposed to the CS in the absence of shock, which normally results in a decrease in fear expression. These responses are adaptive, with the animal first learning that a cue signals threat, and then that the threat is no longer relevant. However, extinction learning can be disrupted by various biological and environmental factors (Singewald & Holmes, 2019). Understanding the mechanisms of fear extinction, especially in the case where extinction is impaired, can help us to understand disorders such anxiety or PTSD, where fearful or traumatic memories are persistent and intrusive (Flores et al., 2018; Singewald & Holmes, 2019).

There is rapidly growing evidence that, as with anxiety disorders, there are sex differences in fear conditioning and extinction (Day & Stevenson, 2019; Velasco, Florido, Milad, & Andero, 2019). However, conflicting findings have been reported. For example, some studies showed facilitated conditioning in males compared to females (Gupta et al., 2001; Wiltgen et al., 2001; Baran, Armstrong, Niren, & Conrad, 2010; Kosten, Miserendino, Bombace, Lee, & Kim, 2005), while quite a number observed no sex differences (Wiltgen et al., 2005), and at least one other showed a conditioning deficit in males (Chen et al., 2014). Regarding extinction, some studies showed that females show impaired extinction compared to males (Baran et al., 2009), while others showed equivalent performance (Milad et al., 2009). While sex differences in behavioral measures of fear during extinction are not always reported in humans, differences in neural activity within fear circuitry during extinction have been observed (Lebron-Milad et al., 2012). To date there have been no studies that investigate sex difference in fear conditioning or extinction in adolescents.

The discrepancies described above may be due to the effect of estrous phase and related cycling gonadal hormones on conditioned fear (Maeng et al., 2017; Wegerer et al., 2014; Zeidan et al., 2011). For adult female humans and rodents, extinction facilitation is observed where natural levels of estrogen are high (Wegerer et al., 2014; Zeidan et al., 2011). This protective effect of estradiol is interesting, given that anxiety is less prevalent in males (Baxter et al., 2012) where levels of estradiol are low (Gibbs, 2005). It may be that gonadal hormones generally promote extinction. For example, increasing serum testosterone levels (via an acute injection of leuprolide acetate, a gonadotropin-releasing hormone receptor agonist) enhances extinction in adult male rats (Maeng et al., 2017).

Expression of fear conditioning and extinction is also age-dependent (Ganella & Kim, 2014). For example, while renewal and reinstatement of extinguished fear are robust in adult rats (Ganella & Kim, 2014; Singewald & Holmes, 2019), they are impaired in juvenile male but not female rats (Park et al., 2017). On the other hand, adolescents show impaired extinction compared to adults, demonstrated in male rodents (Kim et al., 2011; Pattwell et al., 2012; Zbukvic et al., 2017; Baker & Richardson, 2015), and male and female humans (Ganella et al., 2018; Pattwell et al., 2012). This extinction deficit is thought to be linked adolescent vulnerability towards anxiety disorders (Baker, Bisby, & Richardson, 2016). Surprisingly, there has been no research to date that examined sex differences and/or the mediating effect of gonadal hormones on extinction during adolescence.

Adolescence is a time where the onset of puberty leads to large fluctuations in gonadal hormones that were not present earlier in life (Abreu and Kaiser, 2016). This is important because hormonal fluctuations increase vulnerability to anxiety and other affective disorders (Brummelte and Galea, 2010) and indeed menarche has been linked with heightened risk of anxiety in females (Patton et al., 1996). It is possible that the effects of hormones on behavior is different during this stage of development when compared to adulthood.

The aim of this study, therefore, was to investigate sex differences in Pavlovian fear conditioning and extinction in adolescent rats, and whether estrous cycling mediates such difference. In experiment 1, male and female rats underwent Pavlovian fear conditioning beginning on Postnatal day (P)35. We hypothesized that females would show delayed extinction learning and impaired extinction recall compared to males. Similar to adult animals, extinction impairments in female adolescents may also be the most pronounced during estrus, when estradiol level is the lowest in rodents (Smith et al., 1975; Staley and Scharfman, 2005). In experiment 2, we tested the hypothesis that gonadal hormones generally promote extinction. We manipulated gonadal hormone release by performing ovariectomy (OVX) or orchidectomy (ORX) surgery in P21 male and female rats, and then trained rats in Pavlovian fear conditioning at P35. We also verified plasma estradiol levels at each estrus phase in these rats. The hypothesis would be supported if OVX or ORX cause impaired extinction and extinction recall.

## 2. Materials and methods

### 2.1. Subjects and surgery

Male and female Sprague Dawley rats were bred in-house at the Florey Institute of Neuroscience and Mental Health, Melbourne, Australia. Rats were weaned at P21 and housed for the remainder of experimental procedures with same-sex littermates in groups of 5-6 in individually ventilated cages under a 12/12 h cycle (lights on at 07:00) with food and water available *ad libitum*.

For experiment 2, all rats received ORX or OVX surgery before weaning on the same day.

Rats were anaesthetized with an isoflurane/oxygen mixture (5% for induction, then 1.5-2% for maintenance). For male rats, an incision was made on the scrotum, and testes were ligated and removed. For female rats, two incisions were made on the dorsal side above the location of the ovaries to remove the ovaries. Sham surgeries were identical to ORX and OVX surgeries except testes or ovaries were left intact.

All rats were P35 (±1) on the first day of behavioral training, and were handled three times prior. All procedures were approved by the Animal Care and Ethics Committee at the Florey Institute of Neuroscience and Mental Health in accordance with the guidelines for animal use set out in the Australian code of practice for the care and use of animals for scientific purposes (8^th^ edition, 2013).

### 2.2. Apparatus

All behavior occurred in *Contextual Near Infra-red (NIR) Fear Conditioning System* and *Video Freeze* system (Med Associates, VT, USA). The dimensions of the chambers and the grid floor were as described previously (Zbukvic et al., 2017). The chambers contained one of two different contexts, located in two individual rooms. Briefly, the first context had houselights on, round stickers on the back wall, woodchip bedding beneath the grid floor, and was cleaned with soap containing a mild eucalyptus odor. The second context had curved walls, a tray of paper towel placed beneath the grid floor and was cleaned with 80% v/v ethanol. These two contexts were fully counterbalanced to serve as conditioning and extinction contexts. During extinction and test sessions, a Perspex sheet was placed across the grid floor to further differentiate the contexts.

### 2.3. Procedures

#### Conditioning

Rats were placed in the novel chambers and after 2-minutes, the CS (tone, 5000Hz, 80 dB) was presented for 10 seconds, co-terminating with the 1 second US (foot-shock; 0.6 mA). Rats received 6 CS-US pairings [inter-trial interval (ITI; 85-135 seconds)].

#### Extinction

The next day, rats were placed in a different context to fear conditioning. After 2 minutes, the 10 second CS was presented alone 60 times (10 second ITI).

#### Test

The day after extinction, rats were placed back in the extinction context. After 2 minutes, the 10 second CS was presented alone 5 times (10 second ITI).

In all behavioral sessions, freezing was calculated via NIR video tracking equipment and computer software (Video Freeze, Med Associates, VT, USA). Freezing was defined by a motion threshold of <50 pixel changes per frame for a minimum period of 30 frames (1 second), a calibration previously shown to have high concordance with manual scoring in adolescent rats (Ganella et al., 2017b).

### 2.4. Estrous cycle monitoring

For experiment 1, vaginal lavages were taken from female rats 1 hour after each behavioral session, and 1 day after the extinction recall test day. For experiment 2, lavages were also performed immediately before collection of trunk blood for a sample of females in each group. A pipette tip containing 20uL of saline was inserted ~1-3 mm into the vagina, unless the vagina was unopened. The area was flushed two or three times, then the fluid collected was transferred to a microscope slide and air dried before staining with a 4% (v/v) methylene blue solution for 15 minutes. Stained slides were rinsed twice with distilled water and air dried before observation under a light microscope (10x magnification, Olympus BH-2). Slides were blinded and cross-checked by two independent researchers. They were categorized according to the 4 discernible phases of the rat estrous cycle: proestrus, estrus, metestrus and diestrus. The morphology of cells and their relative cellular proportions were used to determine the phase of estrous (Cora, Kooistra, & Travlos, 2015 - Figure 1a). For experiment 1, rats in diestrus and metestrus were pooled together for statistical analyses because only a few rats were in the metestrus phase, a method adopted in other studies (Hecht et al., 1999) due to the relatively short length of metestrus compared to other phases and similar rising levels of estrogen through these phases (Westwood, 2008).

**Figure 1.**
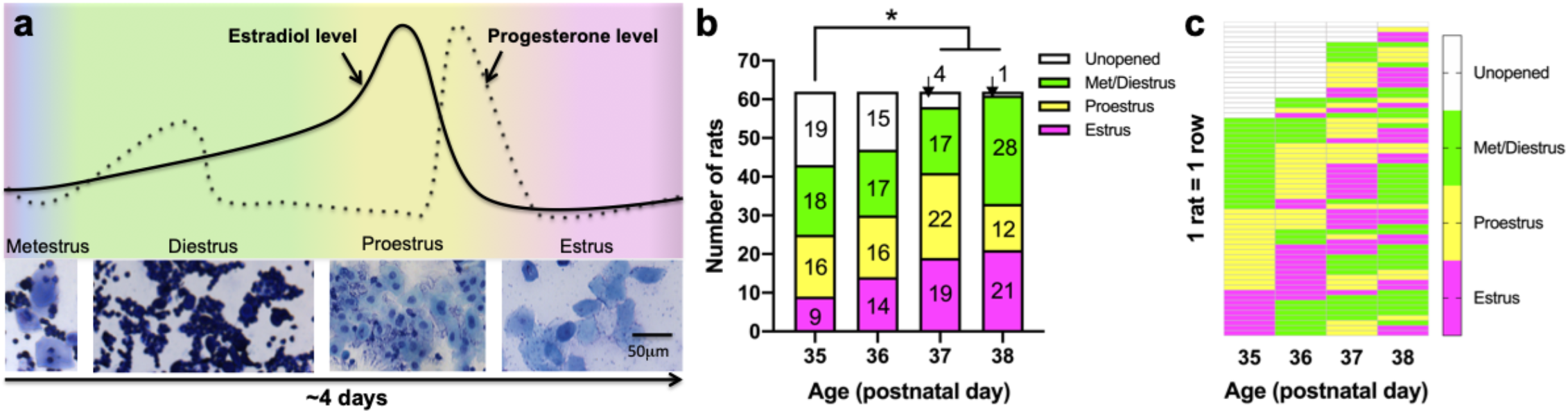
Estrous phase monitoring and maturation. **(a)** Representative vaginal cytology smears for each phase of the rat estrous cycle in the present study. Briefly, proestrus is characterized by small, round mononucleated cells; estrus by anucleated keratinised epithelial cells; metestrus by anucleated keratinized epithelial cells with interspersed leukocytes that can be differentiated from other cells by their small size, and diestrus which is predominated by leukocytes (Cora et al., 2015). Depicted estradiol and progesterone levels are based on levels measured every 4 hours in Sprague-Dawley rats as reported in (Smith et al., 1975). The arrow underneath indicates passage of time. **(b)** Number of rats at each phase of estrous cycle across experiment 1 (For P37 and P38, unopened n/group is indicated by arrows above the column). Numbers of rats in each phase significantly differed between P35 and P37/P38, with the decrease in the number of rats with unopened vaginas (*ps < 0.05). Total female n = 62. **(c)** Estrous phase for each rat (1 rat = 1 row) on each day P35-P38.

### 2.5. Blood collection and ELISA

In experiment 2, female rats were killed by cervical dislocation for blood collection via atrial puncture 24 hours following test. Blood from a random selection of male rats (6 sham and 6 ORX) was collected the same way in order to confirm that the surgery was effective in reducing testosterone levels in circulating blood to negligible levels. Whole blood was collected in lavender EDTA Vacutainers (Becton Dickinson, Scoresby, VIC, Australia) and plasma was isolated by cold centrifugation for 10 minutes at 1500G. Plasma was stored at −20C until further use. Samples were analyzed using commercially available enzyme-linked immunosorbent assay (ELISA) kits in triplicate for estradiol (#582251, Caymen Chemical, Ann Arbor, MI, USA), progesterone (#582601, Caymen Chemical, Ann Arbor, MI, ISA) and testosterone (LS-F39213, LifeSpan Biosciences, Seattle, WA, USA). As reported in previous studies (Feng et al., 2007; Lv et al., 2016), we followed manufacturer’s instructions. To summarize, samples and standards were loaded to the 96-well plate in triplicate for each essay. Two wells were designated each for blanks, total activity, non-specific binding and maximum binding. Estradiol and progesterone plates were read at a wavelength of 420 nm, and testosterone plate was read at a wavelength of 450nm. The plasma concentrations of each hormone were determined by comparison to standard curves. The resulting assays had a 20-80% B/B_0_ detection range of 9.9-603.8pg/ml for estradiol, 19.3-1862.3 pg/ml for progesterone and a range of 0.313-20ng /ml for testosterone.

### 2.6. Data analyses

All analyses were carried out using SPSS (IBM Corp., New York, NY, USA). Differences in the number of rats in each phase of estrous were analyzed using Chi-Square. Percent freezing from all training sessions was subjected to analysis of variance (ANOVA) or repeated-measures (RM) ANOVA, followed by post-hoc Tukey HSD multiple comparisons or Bonferroni-corrected post-hoc tests wherever appropriate. For analysis of CS-elicited freezing within-session conditioning, the first 9 seconds of each CS-US trial was used to discount the effects of shock on movement. For analysis of CS-elicited freezing within-session extinction, data were collapsed into 12 blocks of 5 CS trials, and for test, 1 block of 5 CS trials. For all experiments, no differences were observed in baseline freezing, which is reported in figures.

## 3. Results

### 3.1. Experiment 1: The relationship between natural estrous cycling and fear conditioning/extinction

In experiment 1 estrous phase was determined on P35, P36, P37, and P38 with vaginal lavages then cytology under a light microscope (Figure 1a), unless the vagina was unopened. As expected, the number of rats in each phase per day significantly differed (Figure 1b), *χ*^2^ (9,62) = 35.74, p < 0.05. Post-hoc tests confirmed that the number of rats in each phase at P35 was significantly different from P37 and P38 (ps < 0.05), driven by the decreasing number of female rats with unopened vaginas (confirming that these rats were undergoing puberty). Figure 1c shows the estrous stage for individual rats on each day. While estrous cycle is irregular for some of these animals, this is not unusual for first cycles following menarche (Laffan, Posobiec, Uhl, & Vidal, 2018).

For behavior, three different sets of analyses were carried out. Specifically, female rats were grouped according to estrous phase/unopened vagina on conditioning day (Figure 2a), extinction day (Figure 2b), and test day (Figure 2c). For each of these groupings, behavioral differences were tested for each day. Males were included as a fifth group.

**Figure 2.**
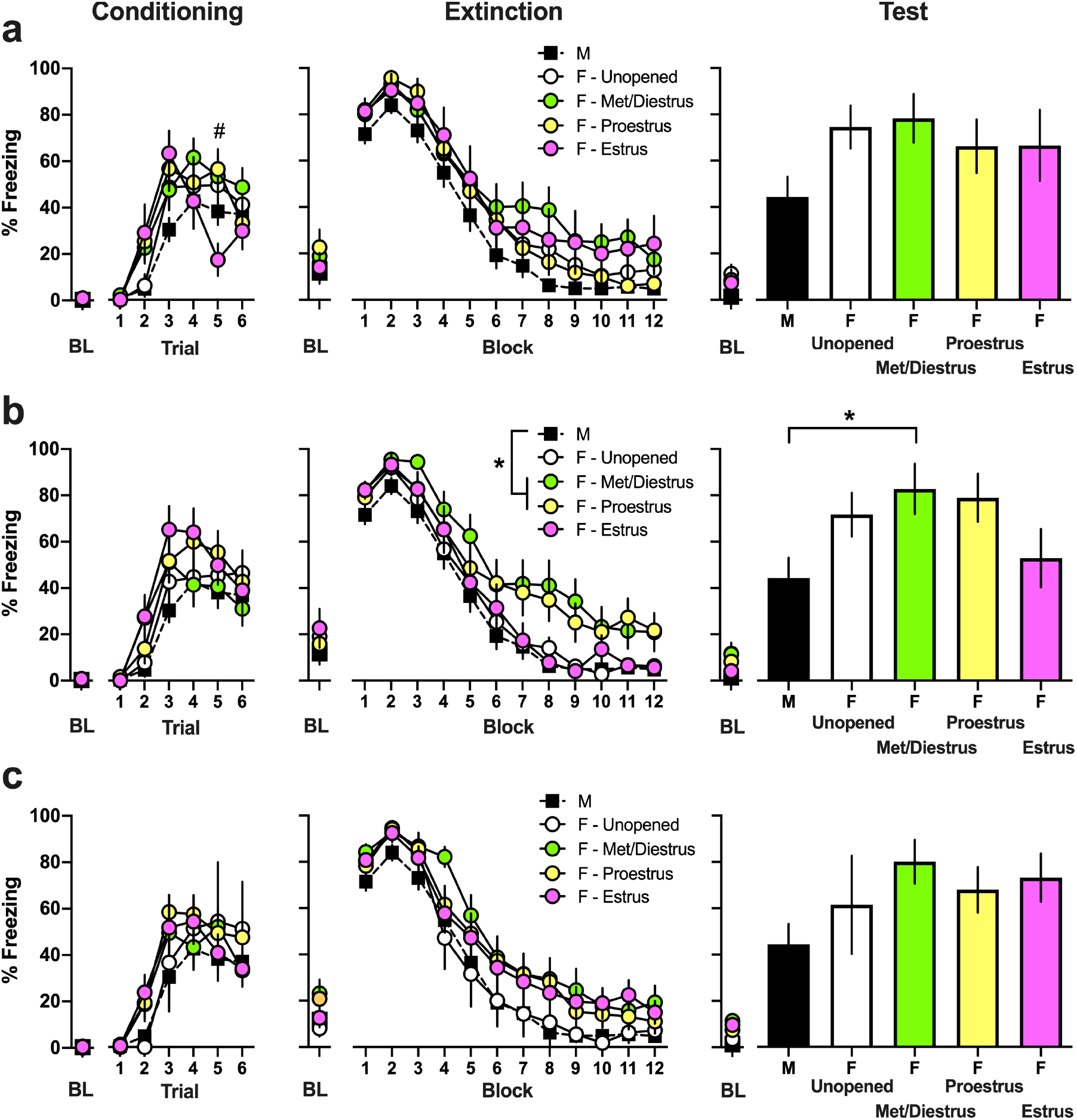
Baseline and CS-elicited freezing in experiment 1. Baseline freezing was not affected by estrous phase on any day. Males (M), Females (F). **(a)** When females were divided into estrous phase at conditioning, there was a significant estrous phase x conditioning trial interaction with females in estrus freezing less than females in met/diestrus or proestrus during trial 5 only (#ps < 0.05). Extinction acquisition or recall was unaffected by conditioning estrous phase. M (n = 26); F – Unopened (n = 19); F – Met/Diestrus (n = 18); F – Proestrus (n = 16); F – Estrus (n = 9). **(b)** When females were divided into estrous phase at extinction, there were no differences during conditioning the previous day. During extinction there was a main effect of estrous phase, with post hoc tests indicating that females in met/diestrus or proestrus froze more than males overall (*ps < 0.05). At test, there was a main effect for extinction estrous phase, with post-hoc tests showing that females in met/diestrus froze more than males (*p < 0.05). M (n = 26); F -Unopened (n = 15); F – Met/Diestrus (n = 17); F – Proestrus (n = 16); F – Estrus (n = 14). **(c)** Estrous phase at test did not affect prior conditioning or extinction, as well as test. M (n = 26); F – Unopened (n = 4); F – Met/Diestrus (n = 17); F – Proestrus (n = 22); F – Estrus (n = 19).

When grouped according to estrous phase on conditioning day (Figure 2a), RM ANOVA of CS-elicited freezing across conditioning trials revealed a significant effect of Trial (F(5,415) = 42.10, p < 0.05), but no effect of conditioning Group (p > 0.05). The Trial x Group interaction was significant (F(20,415) = 1.69, p < 0.05), and post-hoc tests showed that this was due to females in estrus freezing less than females in met/diestrus or proestrus during trial 5 only (there were no other group differences). RM ANOVA of freezing across extinction revealed a main effect for Block (F(11,913) = 171.10, p < 0.05), however there were no effects of Group, and no interaction (ps < 0.05). ANOVA showed that there was no effect of Group on freezing at test (p > 0.05).

When the females were grouped according to estrous phase on extinction day (Figure 2b), RM ANOVA of conditioning trials revealed a significant effect of Trial (F(5,415) = 47.36, p < 0.05), but no effect of Group, nor an interaction (ps > 0.05). For extinction there was significant effects of Block (F (11,913) = 190.64, p < 0.05) and Group (F (4,83) = 4.487, p < 0.05). Post hoc tests showed that females in met/diestrus or proestrus froze more than males (ps < 0.05), with no other group differences (ps > 0.05). There was no Group x Block interaction (p > 0.05). At recall test, ANOVA showed a main effect for Estrous (F(4,83) = 2.97, p < 0.05), and post-hoc tests revealed that females in met/diestrus froze more than males (p < 0.05).

When females were grouped according to estrous phase on test day (Figure 2c), RM ANOVA across conditioning revealed an effect of Trial (F(5,83) = 31.96, p < 0.05) and no other effects (ps > 0.05). Across extinction there was an effect of Block (F(11,913) = 128.379, p < 0.05) and no other effects (ps > 0.05). One-way ANOVA at test revealed no effect of test Group (p > 0.05).

### 3.2. Experiment 2: The effect of gonadectomy surgery prior to conditioning on extinction

Experiment 1 showed that females in met/diestrus or proestrus at extinction froze more than males during extinction overall. Female rats in met/diestrus at extinction also displayed impaired extinction recall when tested the following day. Since met/diestrus is associated with increasing, and proestrus with highest estradiol level (Figure 1a; Staley et al. 2005; Smith et al. 1975), experiment 2 investigated whether changing gonadal hormones levels played a causal role in CS-elicited freezing by performing gonadectomy at P21 and behavioral training beginning at P35.

RM ANOVA of conditioning revealed a main effect of Trial (F(5,255) = 34.28, p < 0.001), and no other effects (ps > 0.05). At extinction, there was a significant effect of Block (F(11,572) = 48.24, p < 0.001), Block x Gonadectomy interaction (F(11,572) = 3.243, p < 0.05), Block x Sex x Gonadectomy interaction (F(1,52) = 4.58, p < 0.05), and no other effects (ps > 0.05). The significant 3-way interaction was followed up by a MANOVA for each block, which revealed significant Sex x Gonadectomy interaction at blocks 4, 5 and 6 (Fs = 6.50 - 8.59, ps < 0.05). These results indicate that OVX females showed accelerated, and ORX males showed delayed, extinction acquisition in the first half of extinction training (Figure 3). At test, 2-way ANOVA revealed no main effects for Sex or Gonadectomy (ps > 0.05), however, there was a significant Sex x Gonadectomy interaction (F(1,52) = 5.078, p < 0.05). Post-hoc tests revealed that OVX rats showed reduced freezing compared with female-sham and ORX groups (ps < 0.05).

**Figure 3:**
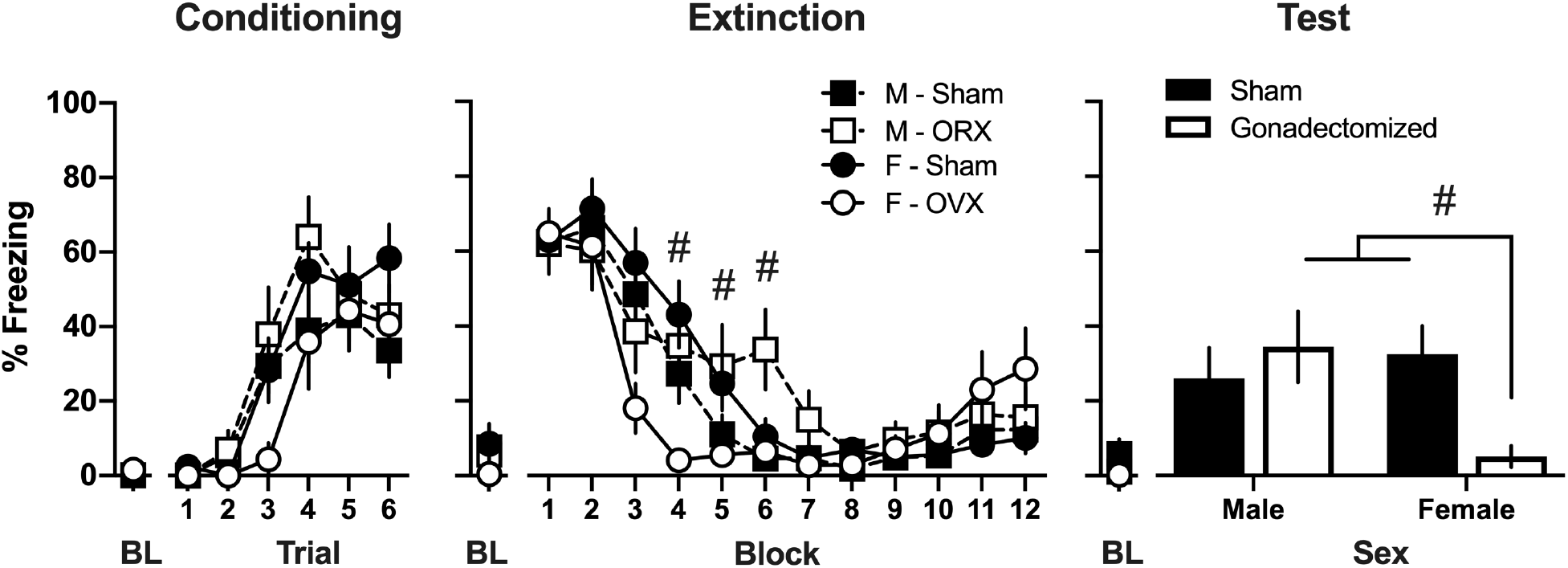
Baseline and CS-elicited freezing in experiment 2. Males (M), Females (F), orchidectomy (ORX), ovariectomy (OVX). While conditioning freezing levels were unaffected by gonadectomy, female OVX rats showed facilitated extinction acquisition and improved extinction recall, while male ORX showed delayed extinction. (#Sex x Gonadectomy interaction on Block 4, 5, 6 and retrieval, ps < 0.05). M - Sham (n = 16); F - Sham (n = 17); M - ORX (n = 12); F - OVX (n = 11).

Due to technical issues, blood samples were lost from 6 sham and 6 OVX females, and 6 ORX males. For remaining female rats, levels of serum estradiol and progesterone were consistent with estrus phase measured immediately before blood collection, and OVX females had negligible circulating estradiol (Figure 4a) and progesterone (Figure 4b) when compared to Sham females (estradiol: t(14) = 3.78, p < 0.001, progesterone; t(14) = 8.88, p < 0.001). When females were divided by estrous phase (only applicable for shams, as all OVX remained immature), one-way ANOVA confirmed that there was a significant difference between groups for estradiol and progesterone [F(1,15) = 46.79 for estrogen; 31.24 for progesterone, ps < 0.05]. Post hoc tests confirmed that hormones were lower in OVX group regardless of phase (all ps < 0.05). Since the purpose of experiment 2 was not to examine effect of phase, power in collected sample size was insufficient to make inferences regarding hormone levels between phases.

**Figure 4:**
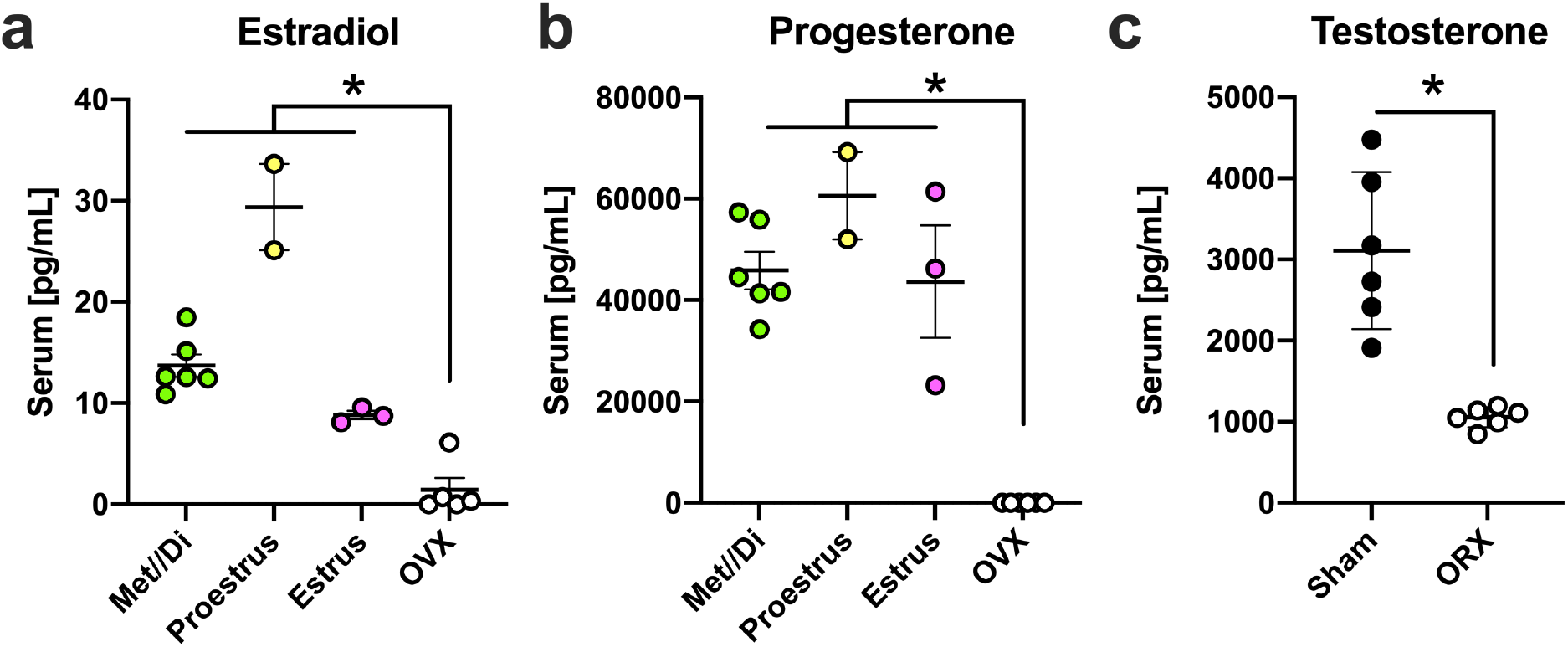
Serum **(a)** estradiol and **(b)** progesterone levels of female rats (Sham (n = 11); OVX (n = 5)), and **(c)** testosterone levels for males rats (Sham (n = 6); ORX (n = 6)), blood taken the day after test. Both Estradiol and Progesterone levels were less in the OVX compared to sham females, regardless of phase. Likewise, testosterone levels were less in ORX compared to sham males (*p < 0.05).

In male rats, we took blood from a random sample of 6 sham male, and all remaining ORX animals (n = 6) to validate the effect of orchidectomy surgery on testosterone. This showed that orchidectomy resulted in circulating testosterone levels (Figure 4c) that were negligible in comparison to sham animals (t(10) = 5.169, p < 0.001).

### 4. Discussion

The present study first showed that adolescent female Sprague-Dawley rats undergo puberty throughout P35-38, with 31% of P35 rats not having begun estrous cycling. This is consistent with previous research (Goldman, Laws, Balchak, Cooper, & Kavlock, 2000; Laffan et al., 2018). For adolescent females, the cycle was somewhat irregular (Figure 1c). This is also consistent with previous research reporting that for many mammalian species, including rats, the first few menstrual cycles are typically irregular (reviewed in Goldman et al., 2000; Laffan et al., 2018). Secondly, estrous phase during extinction significantly affected conditioned freezing in female adolescent rats. Specifically, with females in proestrus or met/diestrus freezing more than males during extinction, and females in met/diestrus freezing more than males at extinction recall test the following day.There was no evidence of a difference between males and females where females were in estrus at extinction, or where females had not yet undergone menarche. In contrast, estrus phase on conditioning day only had an effect on the second-last CS-US trial, and there was no effect on subsequent extinction or test. Estrus phase on test day also had no effects on freezing levels on any behavioral day. Gonadectomy prior to the onset of puberty (i.e., at P21) led to negligible levels of estradiol and progesterone in females, and of testosterone in males. This was associated with a facilitation of extinction and improved extinction recall for female adolescent rats when compared to other groups. Importantly, estrus phase was associated with the lowest level of serum estradiol compared to met/di/proestrus, which is consistent with previous findings in adult rodents, including Sprague-Dawley rats (Smith et al., 1975; Staley and Scharfman, 2005). In contrast to the females, gonadectomy produced delayed extinction in males, although extinction recall was unaffected. This is consistent with the previous finding that gonadotropin receptor agonism enhanced extinction in adult male rats (Maeng et al., 2017).

Together, these results suggest that for adolescent females, unlike for adult females, the presence of high levels of estrogen is associated with impaired extinction. This is consistent with the report that adolescent male rats have low levels of plasma estradiol (Bell, 2018), which may explain the lack of difference between males and females with unopened vaginas or in estrus. In adult rats, it was reported that extinction is facilitated in phases where there are high levels of serum estradiol, although it is puzzling why such phase included estrus (Milad et al., 2009; Zeidan et al., 2011). Notably, those studies did not explicitly measure serum estradiol levels in rats (Milad et al., 2009; Zeidan et al., 2011). We found that rats in met/di estrus (moderate and increasing levels of estradiol) and proestrus (highest levels of estradiol - Figure 1a) (Goldman et al., 2000; Nilsson et al., 2015; Smith, Freeman, & Neill, 1975; Westwood, 2008; Zenclussen, Casalis, Jensen, Woidacki, & Zenclussen, 2014), exhibit increased freezing across extinction and impaired extinction recall the next day (Figure 2b) when compared to males, immature females, or females in estrus (lowest levels of estradiol - Figure 1a; Staley et al. 2005, Smith et al. 1975). Further, females who had undergone OVX surgery prior to the onset of menarche, and consequently had negligible blood estradiol levels (Figure 4b), exhibited decreased freezing across extinction and improved extinction retrieval when compared with intact females (Figure 3a). On the other hand, gonadectomy prior to the onset of menarche resulted in extinction impairment in adolescent males. Again, this is different to studies where extinction was tested in adulthood, which show that orchidectomy (whether pre or post puberty) had no effect on extinction of conditioned fear (McDermott, Liu, & Schrader, 2012). However, increased circulating testosterone was associated with enhanced extinction recall in young adult male rats (Maeng et al., 2017), and testosterone enhanced retention of extinction of a conditioned taste aversion in mice during late adolescence (gonadectomy and testosterone supplementation at P35, behavioral procedures at P49) (Suzuki, Eda-Fujiwara, Satoh, Saito, & Miyamoto, 2013). Those studies strongly suggest that testosterone facilitates extinction in adult male rats, which is consistent with our findings in adolescents.

The conflicting findings regarding the relationship between estrus phase and extinction in females are most likely due to the difference in age in experimental subjects (adolescent in the current study versus adult in previous studies). This age-dependent difference may reflect differences in estrogen receptor (ER) distribution in the brain in adolescence. There are two classical ERs found in the brain – ERα and ERβ (Hara et al., 2015). Although both are important for cognitive function (Hara et al., 2015), binding to each receptor subtype has different effects on behavior (Borrow and Handa, 2017). ERα is associated with anxiogenesis, because its knockdown via ShRNA produced a decrease in anxiety (Spiteri et al., 2010). In contrast, ERβ is associated with an anxiolytic effect, because ERβ knockout mice show increased anxiety-like behavior (Imwalle et al., 2005). In adult animals, estradiol effects on extinction are dependent on binding to ERβ and not ERα (Chang et al., 2009; Zeidan et al., 2011). Notably, the In the cerebral cortex there is a marked decline in ER density leading up to puberty, accompanied by a decrease in ERα mRNA (Wilson et al., 2011). Therefore, alterations in the influence of estradiol levels on extinction in early adolescence vs adulthood might be due increased ERα to ERβ density ratio, although specific changes in densities of ERα or ERβ across adolescence are poorly understood and further research would be required to test this hypothesis.

The neural circuitry that controls fear conditioning and extinction has been well characterized in both adult humans and rodents, with prefrontal cortex (PFC), amygdala, and hippocampus as central structures (Maren, Phan, & Liberzon, 2013). During adolescence, however, there are substantial changes in brain structure, connectivity, and function, including in these regions (Baker et al., 2016; Ganella, Barendse, Kim, & Whittle, 2017; Gennatas et al., 2017; Kim, Perry, Ganella, & Madsen, 2017; Kundu et al., 2018). Specifically, it seems that during adolescence there is decreased recruitment of regions that mediate fear extinction, such as the ventromedial PFC and basolateral amgydala, and increased recruitment of regions that promote fear, such as the central amygdala (Baker et al., 2016). For example, when conditioning and extinction occur during adolescence, levels of phosphorylated mitogen-activated protein kinase (pMAPK) is lower compared to adult rats in the mPFC and basolateral amygdala (Baker & Richardson, 2015; Kim, Li, & Richardson, 2011). In humans, BOLD signals are similarly reduced during fear extinction in adolescents when compared with adults, and this decreased recruitment correlated with extinction impairments (Ganella, Drummond, Ganella, Whittle, & Kim, 2018). Furthermore, adolescent humans showed decreased vmPFCà amygdala recruitment during extinction when compared to adults (Ganella et al., 2017a).

Importantly, estrogen receptors are expressed throughout this circuitry (Ostlund, Keller, & Hurd, 2003), and injection of estradiol has been shown to affect activity within this circuitry. For example, in adult rats undergoing metestrus during extinction, an injection of estradiol immediately after extinction decreased amygdala activity while increasing vmPFC activity measured by the immediate early-gene c-Fos (Zeidan et al., 2011). Similarly, high estrogen was associated with increased activity in the vmPFC, and a positive correlation between vmPFC activity and estrogen levels was observed in female humans (Zeidan et al., 2011). Although similar experiments have not been done in adolescent subjects, it is possible that age-dependent altered connectivity and function of these regions may mediate the different effect of gonadal hormones in adolescence compared to adulthood.

Interestingly, gonadal hormones also cause changes to dopamine innervation of the mPFC. For example, gonadectomy in males increased PFC levels of the dopamine precursor tyrosine hydroxylase (Adler, Vescovo, Robinson, & Kritzer, 1999; Aubele, Kaufman, Montalmant, & Kritzer, 2008; Aubele & Kritzer, 2011), while estrogens enhanced dopamine signaling in this region (Madularu et al., 2015). Dopamine levels in the prefrontal cortex is critical for extinction of conditioned fear in adult animals, and in adolescent deficits in extinction (in male rats) have been linked to changes in ratio of dopamine 1 to dopamine 2 (D1R:D2R), in particular in the medial PFC (Zbukvic & Kim, 2018). This ratio changes across development in a sex-specific way (Cullity, Madsen, Perry, & Kim, 2019) and in response to experience (Zbukvic, Park, Ganella, Lawrence, & Kim, 2017). In fact it is becoming increasingly evident that maturation of the PFC, and in particular changes to dopamine fiber innervation and relative receptor expression, results in divergent mechanisms of cue extinction between adolescents and adults (Zbukvic et al. 2016; Zbukvic & Kim, 2018). Therefore it is possible that the discrepancy between current findings in adolescents and previous findings in adults may be mediated through age dependent changes in dopamine system.

Although our findings do show that extinction is dependent on estrous phase, and that it is affected by pre-pubertal gonadectomy in a sex-dependent manner, we cannot rule out the involvement of factors other than estradiol. In this regard there are two important things worth bearing in mind. First, recent evidence shows that estradiol is still present in the brains of OVX females where circulating hormone levels were negligible. This provides evidence for local production of estradiol, at least in the adult brain (Li and Gibbs, 2019), and hence effects on behavior cannot solely be explained by physiological plasma levels, and estradiol produced by the ovaries. Nevertheless, this does not exclude the possibility that the current results derive from the presence of serum estradiol. Brain levels of estrogen were further influenced by systemic injection of estrogen in a region-dependent manner (Li and Gibbs, 2019). Indeed, given that some of the regions where local production and control of estradiol levels was reported in adult OVX females (e.g. the frontal cortex) are still developing during adolescence, this may in fact help explain why there was a different effect of serum estradiol levels on behavior in adolescence compared to adulthood.

Second, it is possible that effects observed may be due to progesterone levels (Maeng and Milad, 2015) that also show cyclical fluctuations in menstruating females that are subtly different from estrogen fluctuations (see Figure 1a). Injection of allopregnanolone (a metabolite of progesterone) into the bed nucleus of the stria terminalis of adult female rats enhanced contextual conditioning, but not freezing to a CS (Nagaya et al., 2015). On the other hand, another study showed progesterone treatment in OVX rats had no effect on either CS or contextual fear conditioning, although it did improve other measures of cognitive performance (Frye and Walf, 2008). The discrepant findings may indicate that the ratio of progesterone to estrogen is important, which has been shown to affect anxiety (Hiroi and Neumaier, 2006).

Finally, it is also worth noting that for experiment 1 we assumed that hormone levels at each phase of estrous are equivalent to those seen in adult animals (Nilsson et al., 2015). In fact there is, to our knowledge, no study that has tracked hormone levels at each phase of the estrous cycle in adolescent animals, although there are a number of studies that look at estrous cycling from puberty using cytology to classify phases (see Goldman et al., 2000 for a review). All those studies seem to work under the assumption that estradiol levels are similar in adolescents and adults at the equivalent phase. It may be that the difference between the relationship between estrous phase and extinction in adolescent animals (experiment 1), and that seen in adult animals (Milad, Igoe, Lebron-Milad, & Novales, 2009), may be due to the fact that estradiol (or other hormone levels) are different in adolescents. However, experiment 2 does confirm that removal of these hormones in adolescent females facilitated extinction. Further, in experiment 2 we measured hormone levels in order to confirm that the surgeries effectively removed gonadal hormones. These suggested that levels at each phase were consistent with adult data (see Figure 1a and Figure 4), although since the purpose of this experiment was not to examine the effect of phase there was insufficient power to run statistics on these levels. The best study of this would require increased numbers of rats to be tracked every few hours across days, however this was beyond the scope of the current study. Overall it is clear that further research is required into the effect of ovarian hormones on behavior, and that special attention needs to be paid to the effect in adolescent versus adult subjects.

It is important that we acknowledge the limitations in quantifying estradiol using commercially available immunoassays. There are reports of large variations in specificity and accuracy, particularly at low concentrations, and this is an on-going issue for clinical diagnostics and preclinical research requiring precise quantification (Ketha et al., 2015). There is currently no fully satisfactory method to quantify estradiol concentrations in rodents. A previous comparison of the correlative performance of several commercially available immunoassays to liquid-chromatography tandem mass spectrometry (LC-MS/MS) based assays reported that only two kits produced results that correlated with LC-MS/MS (of which only 1 remains available) (Haisenleder et al., 2011). While there may be concerns regarding the preciseness of the low concentrations calculated in our study, it should not distract readers from appreciating the effects of ovariectomy. That is, all the potentially unreliable low values we observed (below 7 pg/ml) are from the ovariectomised animals (Figure 4a). Ovariectomised animals are expected to have little to none circulating estradiol, and the main findings of our study focus on naturally cycling adolescent females. All of our sham controls have estradiol levels above 8 pg/ml (Figure 4a).

It is widely recognized that anxiety- and trauma-related disorders are more prevalent in females in the human population (Velasco et al., 2019). Mirroring this, females rats generally show impaired extinction of Pavlovian fear when compared to males, but estrogen has a protective effect (i.e. promotes extinction) in both adult female rats and humans (Lebron-Milad & Milad, 2012). Here we provide the first evidence for estrous cycle-mediated difference between female adolescent rats in conditioned fear. Importantly, we provide evidence that the protective effect of estrogen seen in adult females (Lebron-Milad & Milad, 2012) is reversed during this stage of development. The finding that increased estradiol at the onset of puberty may be detrimental for fear extinction provides an ethologically valid model to study female vulnerability to anxiety disorders. Aside from the fact that the inability to inhibit inappropriate fear an important cognitive component of anxiety (Ganella and Kim, 2014; Maren et al., 2013; Singewald & Holmes, 2019), many behavioral strategies for treating anxiety disorders involve an aspect of extinction (Singewald et al., 2015). The findings from this study, together with similar studies in adult animals (Zeidan et al., 2011) suggest that when administering these therapies estrous cycle should be taken into account. However, given the discrepancy between previous adult and current adolescent findings in fear extinction, it is likely that the client’s age should also be considered. Indeed, the present study reinforces the importance of developmental research in order to understand how anxiety disorders emerge and can be treated.

## Acknowledgments

We thank Ms Liubov Lee-Kardashyan for carrying out manual scoring where required for both experiments.

